# Chloroplasts of symbiotic microalgae remain active during bleaching induced by thermal stress in Collodaria (Radiolaria)

**DOI:** 10.1101/263053

**Authors:** Emilie Villar, Vincent Dani, Estelle Bigeard, Tatiana Linhart, Miguel Mendez Sandin, Charles Bachy, Christophe Six, Fabien Lombard, Cécile Sabourault, Fabrice Not

## Abstract

Collodaria (Radiolaria) are important contributors to planktonic communities and biogeochemical processes (*e.g.* the biologic pump) in oligotrophic oceans. Similarly to corals, Collodaria live in symbiosis with dinoflagellate algae, a relationship that is thought to explain partly their ecological success. In the context of global change, the robustness of the symbiotic interaction and potential subsequent bleaching events are worth consideration. In the present study, we compared the ultrastructure morphology, symbiont density, photosynthetic capacities and respiration rates of colonial Collodaria exposed to a range of temperatures corresponding to natural conditions (21°C), moderate (25°C) and high (28°C) thermal stress. We showed that symbiont density immediately decreased when temperature rises to 25°C and the collodaria holobiont metabolic activity increased. When temperature reached 28°C, the collodarian host arrived at a tolerance threshold with a respiration nearly stopped and largely damaged morphological structures. Over the course of the experiment the photosynthetic capacities of remaining symbionts were stable, chloroplasts being the last degraded organelles from the microalgae. These results contribute to a better characterization and understanding of temperature-induced bleaching processes in planktonic photosymbiosis.

## Introduction

Radiolaria, and more specifically Collodaria, have been recently unraveled as key players of oceanic ecosystems (Biard et al., 2016; Guidi et al., 2016). Previously overlooked because of their fragility, the use of non-destructive, *in situ*, imaging tools showed that Collodaria represents around 30% of zooplankton biomass at subsurface in oligotrophic oceans (Biard et al., 2016). Collodaria were also reported to be central actors in plankton community networks related to carbon export to the deep ocean via both primary production and vertical flux (Guidi et al., 2016). Initially described by E. Haeckel towards the end of the 19th century (Haeckel, 1887), Collodaria have been little studied so far, but from few landmark studies conducted on radiolarian morphology and physiology (Anderson, 1983).

Collodaria are unicellular organisms that live either solitary (cell size ~1mm) or form colonies as long as few cm in length. Each colony is formed by hundreds to thousands of cells agglomerated in a gelatinous matrix. All cells, individually represented by a spherical structure called the central capsule, are connected together through cytoplasmic extensions and sometimes, can bear siliceous skeleton in some particular species (Anderson, 1976a,b). Collodaria are mixotrophic organisms, feeding on prey from the surrounding environment but also relying on symbiotic interactions with photosynthetic dinoflagellates. Swanberg (1983) suggested that photosymbiosis provides to Radiolaria the minimum energy to subsist, while their energy and nutrients for growth would come from their heterotrophic regime. Collodaria can consume 4 % of their symbionts each day (Anderson 1983) and a part of the photosynthates are transferred to the host mostly as lipid storage pool (Anderson et al., 1983). Microalgae symbionts isolated from the species *Collozoum spp.* have been recently described as *Brandtodinium nutricula*, a dinoflagellate genus belonging to the Peridiniales order (Probert et al. 2014). Based on genetic barcodes and morphological criteria, it has been shown that *Brandtodinium* is a generalist symbiont similarly to *Symbiodinium spp.* and *Pelagodinium beii*, found in association with corals and Foraminifera, respectively (Decelle et al 2015).

The breakdown of the symbiotic associations in corals, so-called coral bleaching, has been originally described by Glynn (1984), and depicted coral fading as a reaction to environmental constraints, due to the massive loss of symbiotic algae, or alternatively to a decline of their photosynthetic pigment content. Recent projections suggest that more than half of the coral reefs worldwide will experience several bleaching events by year 2100, as a consequence of ocean temperature rise of 1 to 1.8°C (Logan et al., 2014). Coral bleaching has significant deleterious consequences on coral reef ecosystem functioning and as a result, on several countries economy and food supply (Hoegh-Guldberg et al., 2007). Numerous studies are trying to decipher the relationships between corals -or other cnidarians such as sea anemones- and their symbionts upon environmental perturbations. Although bleaching events involve the whole holobiont and all possible interactions between its biotic component (Baird et al., 2009), in cnidarians most studies considered bleaching as being induced by the microalgal symbiont partner release (reviewed in Lesser et al. 2011). One of the proposed mechanism suggests the production of Reactive Oxygen Species (ROS) when symbiont photosynthesis is impaired by heat stress (Lesser et al., 2006). These ROS lead to the death of the host cells by triggering apoptosis, notably through the induction of proteases (Cikala et al, 1999). During the stress and death of the host cells, microalgal symbiotic cells can be either healthy (Baghdasarian and Muscatine, 2000) or damaged (Fujise et al., 2014). In the later case, it is unclear if the symbiont damages are due to the host cell apoptosis or other symbiont related process such as programmed cell death (PCD) such as host autophagic digestion (Dunn et al., 2007; Paxton et al., 2013).

Considering the importance of Collodaria both in terms of abundance and contribution to the biological carbon pump (Biard et al., 2016, Guidi et al., 2016), collodarian bleaching could have a strong impact on oceanic ecosystem functioning. In this study, we performed a controlled thermal stress experiment on Collodaria to investigate plankton photosymbiosis response to elevated temperatures using both morphological and physiological analyses.

## Method

### Experimental design

Collodaria colonies were collected in the bay of Villefranche-sur-Mer (France, 43°41’10” N, 7°18’50” E) using a Regent plankton net (680 μm mesh size), immediately handpicked and transferred into clean beakers filled with 0.2 μm filtered seawater. Colonies were incubated at seawater temperature (21°C) for 3-4 hours to allow self-cleaning of particles embedded in their gelatinous matrix. A total of 300 similar morphotypes were selected and transferred individually into two 17L-Kreisel tank aquarium (JHT-17G, Exotic Aquaculture, Hong-Kong) filled with 0.2 μm filtered seawater and placed in two distinct thermostatic incubator set at 21°C (TC135S, Lovibond Water testing, United Kingdom). Two liters of seawater from the Kreisel tanks were changed on a daily basis. Electroluminescent diode ramps (Rampe led Easyled 6800K°, Aquatlantis, Portugal) provided constant white light at 100 μmol photon m^−2^ s^−1^. Colonies were further exposed to two thermal treatments during 3 days (T0, T1 & T2). In one incubator, the control treatment (CT) remained at 21°C over the course of the experiment, whereas in the other incubator, the stressed treatment (ST), the temperature was incrementally raised everyday 12 hours with sampling from 21°C (T0), to 25°C (T1) and finally to 28°C (T2). In the following text, a “condition” refers to the unique combination of a treatment and a sampling time point (e.g. CT-T0).

### Phylogenetic analysis

RNA sequences were obtained from transcriptome sequencing of specimens collected during the experiment. 18S rDNA gene sequences from the Collodarian host and the dinoflagellate symbionts were retrieved from the transcriptome data by searching for similar HMM profiles (see details in Supplementary Text 1).

### Transmission electron microscopy (TEM)

Collodarian specimens were fixed for 2 hours at room temperature with 2.5% glutaraldehyde in a mix of cacodylate buffer (0.1 M, pH 7.4) / artificial seawater, then washed with 0.1 M cacodylate buffer (pH 7.4) and postfixed with 1% osmium tetroxide in cacodylate buffer containing 1% potassium ferrocyanide. The samples were embedded in Epon resin after 10 minutes dehydration in 90% acetone and 2 baths of 10 minutes in acetone 100%. Ultrathin sections (70-80 nm) of colony parts were cut using a diamond knife mounted on an ultramicrotome (Ultracut S, Leica) and placed on copper TEM grids coated with formvar film. To increase the contrast, the grids were treated with conventional uranyl acetate stain followed by lead citrate. Samples were observed under a JEOL JEM 1400 transmission electron microscope equipped with a CCD camera (Morada, Olympus SIS) at the Centre for Applied Microscopy (Université Côte d’Azur, Nice, France). At least three independent sample observations were conducted for each condition.

### Oxygen measurements

Oxygen consumption was measured using oxygen optodes, as described in Lilley et al. (2014). Two colonies were incubated in 5 mL tubes filled with filtered seawater and equipped with light-sensitive oxygen spots (PreSens Precision Sensing GmbH, Germany). For each condition, measurements were made in 5 replicates. After 30 minutes of dark-acclimation, oxygen was measured using a Fibox 3 optical oxygen meter (PreSens) each 20 minutes during 2 hours. Light (100 μmol photon m^−2^ s^−1^) was then switched on during two hours and oxygen was once again measured every 20 minutes during 2 hours at light exposure.

After calibration, the phase delay measured by the sensor was converted into O_2_ concentration (in μmol L^−1^). Linear regressions between oxygen concentration against time were fitted through all data points per replicate for dark and light measurements separately. Points showing standardized residues <-1.5 and >1.5 were considered as outliers and removed from the dataset to ensure linearity. Slopes of the fitted regression gave individual consumption rates µmol O_2_ L^−1^ h^−1^) and blank slopes were removed from experimental slopes. Respiration rates were estimated as the slope during the dark experiment and gross photosynthesis rates were computed as differences of slopes between light (net photosynthesis) and dark. For normalization, rates were divided by the colony biomass estimated from binocular inspection (see the following morphological measurement methods).

### Chlorophyll fluorescence measurements coupled with microscopy

For each sampling time and treatment, 5 colonies were transferred to cavity slides and maintained with a coverslip to avoid movement. Photosynthetic parameters of individual symbiotic microalgae were measured on three different fields of observation using a Pulse Amplitude Modulated fluorometer coupled to a microscope (MICROSCOPY-PAM; Walz, Effeltrich, Germany), equipped with a 10× objective lens. After the colonies were incubated in dark for at least 5 minutes, the basal level of fluorescence (F_0_) was measured under modulated light (9 μmol photons m^−2^ s^−1^, frequency: 8Hz at 625 nm), and a saturating light pulse (1707 μmol photons m^−2^ s^−1^, with 8×60 ms width at 625 nm) allowed the determination of the maximum fluorescence level (F_M_). The Dark-adapted maximal quantum Yield (F_V_/F_M_) of photosystem II (PSII) was computed as follows:

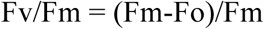

The colonies were then exposed to actinic light (463 μmol photons m^−2^ s^−1^ at 625 nm) during 2 min followed by another saturating light pulse to measure the light-adapted PSII fluorescence yield (F_v_’/F_m_’; Genty et al. 1989). Pilot experiments and light curves were previously performed to determine the optimal duration of the dark incubation and actinic light intensity. The saturating pulse setting were adjusted according to preliminary experiments performed on cultures of *Brandtodinium nutricula* (Supplementary Text 2).

Between 6 and 15 areas of interest corresponding to single symbiotic cells were selected on each picture (see Supplementary Material Figure 1). Outliers were removed when measured values were inconsistent (e.g. F_v_/F_m_=0 or 1) being mostly due to field depth impairments. We obtained between 13 and 45 measurements for each of the 5 replicate colonies per experimental condition. To avoid bias due to these sampling size differences between conditions to compare, the F_V_/F_M_ medians were computed for each microscopic field (3 microscopic fields per colony; 5 replicated colonies per condition).

### Morphological measurements

Images of colonies used for oxygen and PAM fluorescence analyses were acquired under a binocular microscope (Zeiss Stemi SV11 mounted with an Olympus DP21 camera system) and analyzed using the imageJ software (Rasband, 1997; https://imagej.nih.gov/ij/). Stressed colonies at sampling time 2 (ST-T2) were too degraded to manipulate and observe them under the binocular microscope. Therefore, for this specific condition, morphological measurements were performed for one colony and were provided as indicative trends only. Correction factors were applied to normalize measurements in function of the different magnifications used to take each picture. On large field of view, we measured colonies area, width and derived biovolume estimates from the prolate ellipsoid equation described in Biard et al. (2016). We also enumerated the total number of central capsules present on one side of the colonies. On zoomed in fields of view, we measured central capsule area, and central capsule area covered by microalgal symbionts. As previously, we counted the total number of central capsules on one side of the colonies. We also enumerated the symbionts between the central capsules (in the matrix) and measured their areas. As the symbiont cell area was very homogeneous, we derived the number of symbionts per central capsule by dividing their total area by the average area of a symbiont. Symbiont density was thus expressed as the number of symbiont per area. Carbon content was derived from the central capsule counts using a conversion factor of 131 ngC per capsule (Michaels et al., 1995).

### Statistical analysis

For fluorescence and oxygen analysis, a two-way analysis of variance with interaction (ANOVA) followed by Tukey *post hoc* test (for multiple comparisons) was employed and the level of significance was set at p ≤ 0.05, using the following model:

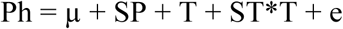

where Ph is the phenotype measured, SP is the sampling time point and T the treatment.

For symbiont density analysis, a T-test was applied to compare regression slopes at each sampling time between the two treatments.

## Results

Colonies were identified as *Collozoum pelagicum* by phylogenetic analysis of 18S rDNA gene sequence (NCBI accession MG907123, Supplementary Material Figure 2) onto a reference tree published by Biard et al. (2015). The sampled specimens belong to the most diverse and abundant collodarian clade C7 from the Sphaerozoidae family which is particularly dominant in coastal biomes and in the western part of the Mediterranean Sea (Biard et al., 2017). We were also able to retrieve 18S rDNA sequences of a dinoflagellate identified as *Brandtodinium nutricula* by phylogenetic placement (NCBI accession MG905637, Supplementary Material Fig. 3), confirming that *C. pelagicum* hosts *B. nutricula* as a symbiotic algae.

### *C. pelagicum* morphological changes upon heat stress

Depending on colony size, an average of 800 (± 600) collodarian cells were agglomerated in a gelatinous matrix, corresponding to an average density of 10 cells per mm^2^. (Supplementary Material Table 1). Cells were distributed in the matrix so as to form a well defined compartment (Fig. 1a). Central capsules, appearing bright under the microscope (Fig. 1a), exhibited a size ranging from 100 to 150 μm. Each central capsule contained a large electron dense droplet, an endoplasm and an ectoplasm (Fig. 1b). Soudan Black staining suggested that this droplet, surrounded by a vacuolar membrane was constituted of fibrous elements and lipids. The endoplasm also contained multiple nuclei that displayed a fibrillar nucleoplasm chord-like structures of chromatin attached to the inner surface of the nuclear membrane (Fig. 1c, d). The endoplasm was delimited from the ectoplasm by a double-layer central capsule membrane and a monolayer vacuole membrane on each side (Fig. 1e).

**Fig. 1:**
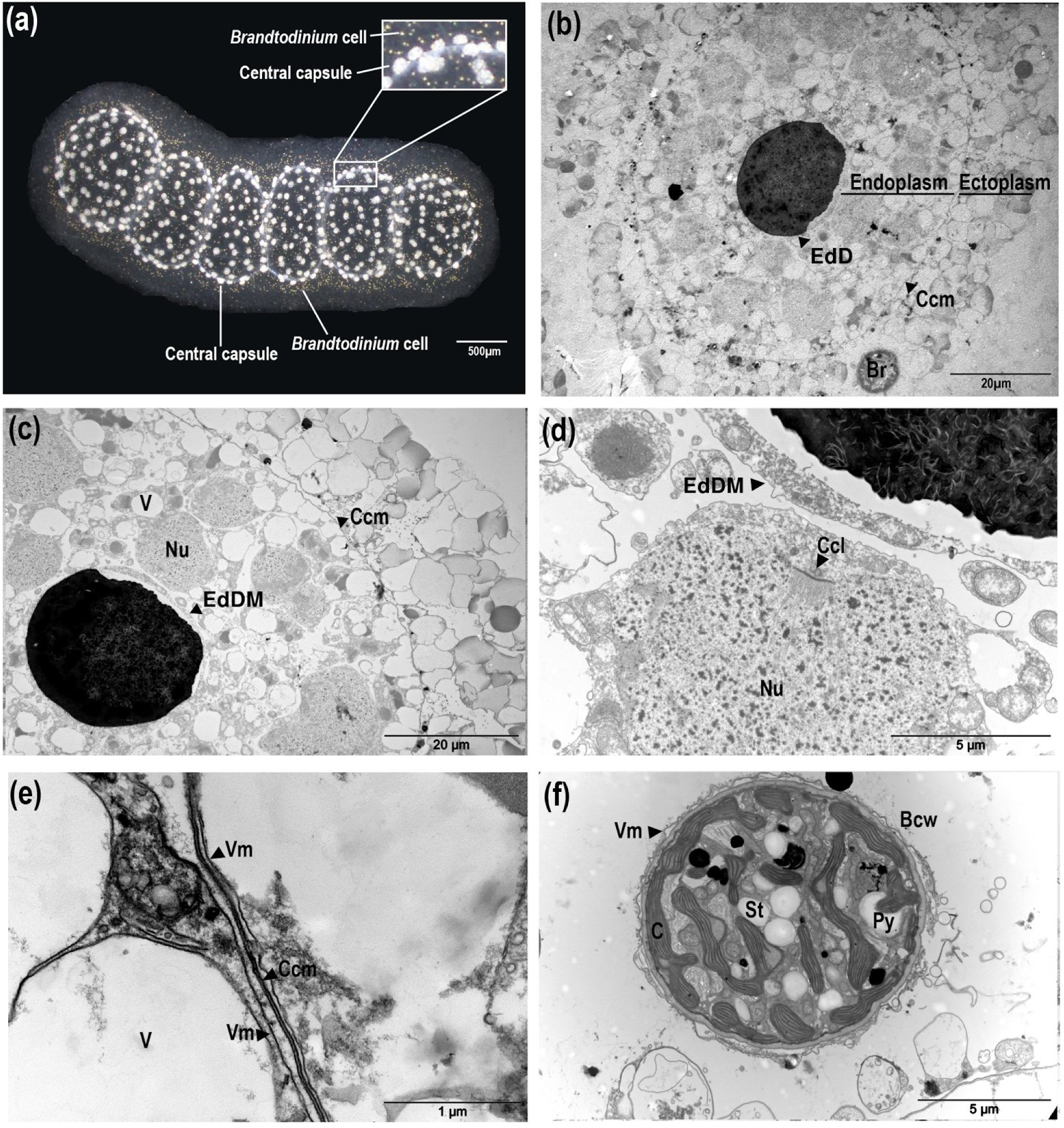
Description of *Collozoumpelagicum* morphology and ultrastructure. *Collozoum* colony morphological aspect by binocular observation (**a**). Observation of the general organization of a central capsule observed by Transmission Electron Microscopy **(b-c).** Focus on nuclei in the endoplasm of the central capsule **(d)**, membranes at the periphery of the central capsule **(e)** and *in hospite Brandtodinium* cells **(f)**, observed by electron microscopy. EdD, Electron-dense Droplet; Ccm, Central capsule membrane; V, Vacuole; Nu, Nuclei; Vm, Vacuolar membrane; EdDM, Electron-dense Droplet Membrane; Ccl, cord-like Chromatin; C, Chloroplast; Bcw, *Brandtodinium* cell wall; Py, Pyrenoid; St, Starch granule.

The dinoflagellate symbionts were always enclosed in cytoplasmic structures, either localized within the gelatinous matrix or closely associated to the central capsules (Fig. 1a, b). Symbiotic *Brandtodinium* exhibited a thick cell wall and no flagella. Symbionts were enclosed within a vacuole and were separated from collodarian ectoplasm by a vacuolar membrane (Fig. 1f). The symbionts contained pyrenoid-bearing lobed chloroplasts, essentially located at the cell periphery, and associated with starch granules. To study responses of *C. pelagicum* to thermal stress, we compared the phenotypes of control colonies (CT at 21°C) to heat stressed colonies (ST) at 3 sampling times (T0:21°C, T1:25°C and T2:28°C). The general morphology of control colonies was preserved during the entire experiment: the polysaccharide matrix surrounded the evenly distributed central capsules and the compartments were still visible (Fig. 2a,b). In heat-stressed conditions, the central capsules from the colonies were still observable but irregularly distributed (Fig. 2c, 2d). The gelatinous matrix was altered at 25°C and completely disorganized at 28°C, while compartments slightly disappeared, revealing large bubbles inside the colonies (Fig. 2d). With respect to ultrastructural features, at T2 for CT colonies (Fig. 2f) and earlier (from T1) in ST colonies (Fig. 2g) ectoplasms were degraded, revealing large spaces around vacuoles. Ectoplasm of ST-T2 colonies was completely shrunk (Fig. 2h). Compared to freshly collected specimens, both control and thermal stress experiments showed highly condensed chromatin, with an increase in electron-dense filaments (Fig. 1c-d, Fig. 2i-l).

**Fig. 2:**
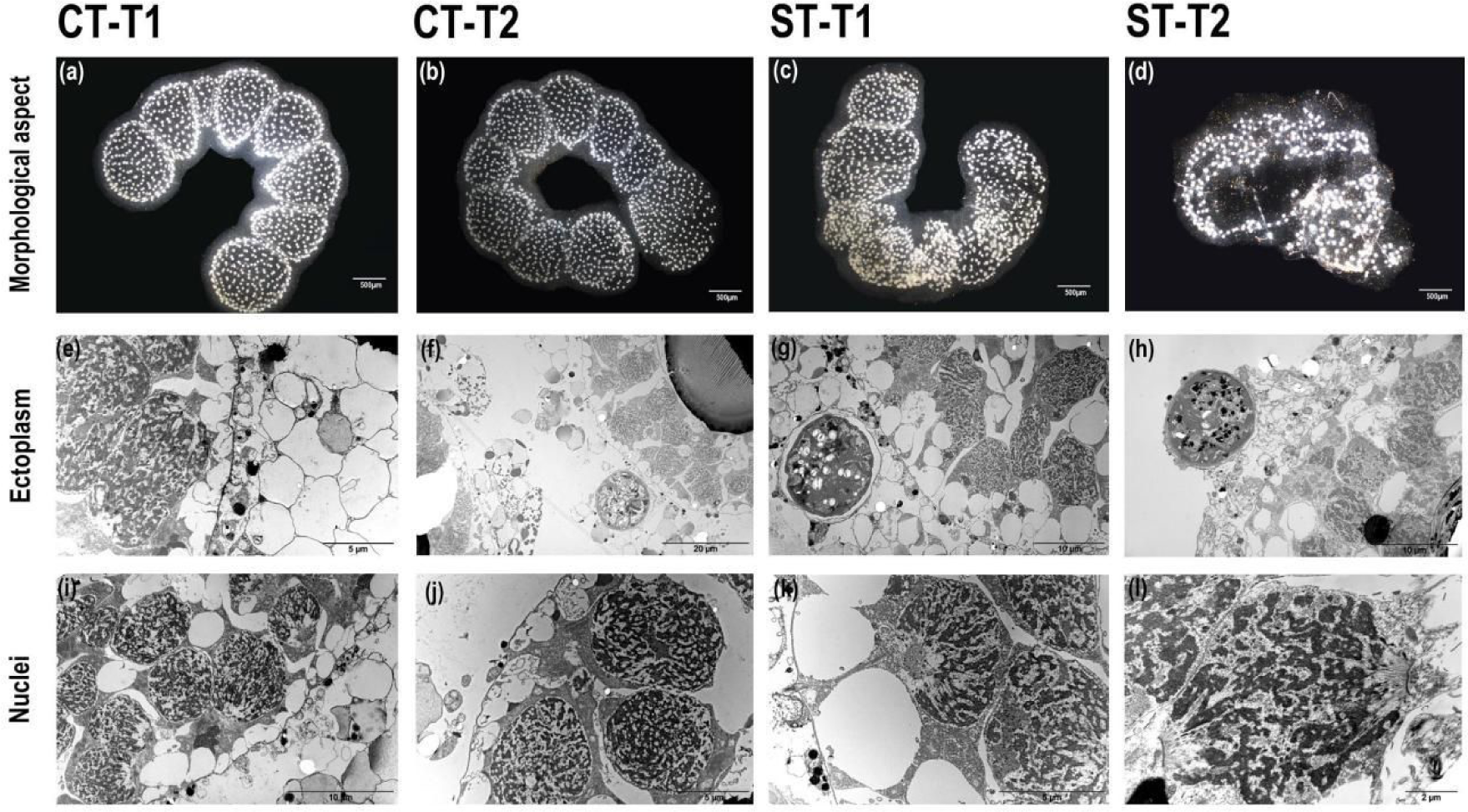
Morphological and ultrastructural changes between control and heat-stressed *Collozoum pelagicum.* Observation of colonies overall morphological aspect at control T1 **(a)** and control T2 **(b)** at 21°C, thermal stress T1 at 25°C **(c)** and thermal stress T2 at 28°C **(d)**. Focus on the ultrastructure of ectoplasm at control T1 **(e)**, control T2 **(f)**, thermal stress T1 **(g)** and thermal stress T2 **(h)**. Focus on the nuclei ultrastructure at control T1 **(i)** and control T2 **(j)**, thermal stress T1 **(k)** and thermal stress T2 **(l)**.

### Physiological impact of heat stress on *C. pelagicum*

At seawater temperature, the respiration rate was estimated to be 3.94 ± 0.99 μL O_2_ mg C^−1^ h^−1^ (Fig. 3, Supplementary Material Table 2). The incubation conditions did not affect significantly the respiration rates in control colonies, with values of 4.58 ± 0.83 μL O_2_ mg C^−1^ h^−1^ at T1 and 4.84 ± 1.25 μL O_2_ ind^−1^ h^−1^ at T2. For heat-stressed colonies, the respiration rates slightly increased at 25°C, reaching 5.82 ± 1.62 μL O_2_ mg C^−1^ h^−1^, and decreased down to 2.94 ± 1.56 μL O_2_ mg C^−1^ h^−1^ at 28°C. The experimental conditions induced a decrease of the average symbiont density from nearly 80 symbionts/mm2 at T0 and T1, down to 59 symbionts/mm^2^ at T2, for control colonies (Fig. 4). The symbiont density decrease occurred significantly faster in heat-stress conditions with as few as 60 symbionts/mm^2^ at T1 (25°C). The regression line slopes between T0 and T1 were significantly different with a stronger drop in symbiont densities in stressed colonies than controls (p-value: 0.05). At T2 (28°C), symbiont density kept decreasing down to 50 symbionts/mm2, but the reaction norm slopes were not significantly different from the control conditions between T1 and T2 (Fig. 4). As described by Anderson (1976a) with electron microscopy in specimens of *C. inerme*, the symbionts were mainly located in the vicinity of the central capsules, while nearly 8% of the symbionts were found in the cytoplasmic extension throughout the gelatinous matrix. During our experiment, the symbiont density varied for the dinoflagellates located nearby the central capsule but remained stable for those in the matrix (Fig. 4).

**Fig. 3:**
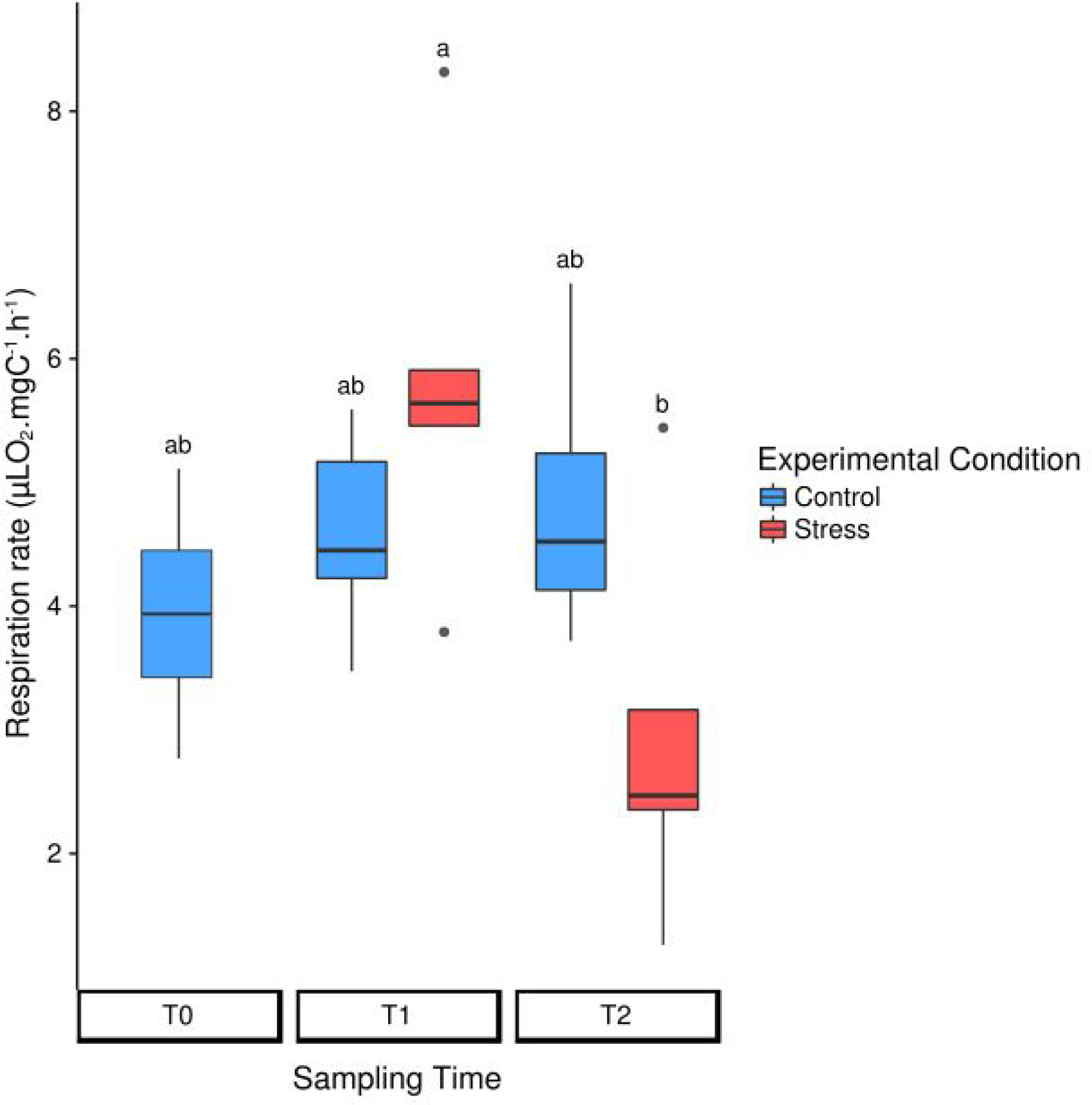
Normalized respiration rates in control (in blue) and heat-stressed (in red) colonies of *Collozoum pelagicum.* The rates are normalized by surface area of the colony. Control colonies were maintained at 21°C, while stressed colonies were sampled at 25°C (T1) and 28°C (T2). 2 way-ANOVA results testing Sampling Time, Treatment and Interaction effects are represented by Tukey post-hoc test associated letters for Interaction effect.

**Fig. 4:**
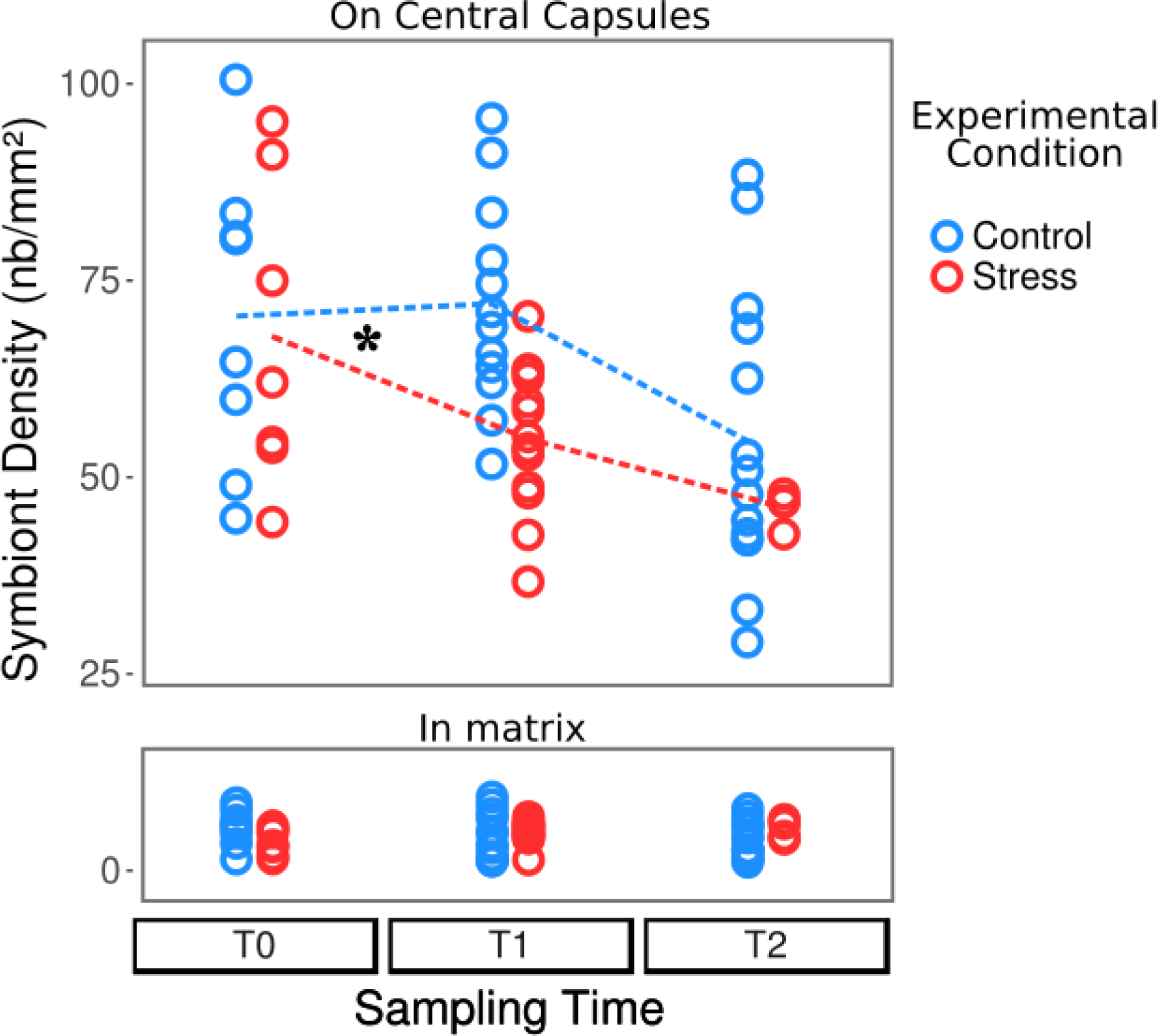
Variations of the symbiont density in control (in blue) and heat-stressed colonies (in red) of *Collozoum pelagicum*. Control colonies were maintained at 21°C, while stressed colonies were sampled at 21°C (T0), 25°C (T1) and 28°C (T2). The upper and bottom parts represent the density of symbionts attached to central capsules and in the gelatinous matrix, respectively. Slopes of reaction norms for symbionts from central capsule (represented as dotted lines) were significantly different between T0 and T1 (p-value <0.001), but not between T1 and T2.

We also performed two different assessments of photosynthetic parameters. First we measured the oxygen production of entire colonies to globally estimate their overall photosynthesis, then we carried out fluorometric measurement coupled to microscopy to evaluate the photosynthetic efficiency of PSII for individual symbiont cells. Both methods showed that the photosynthetic apparatus was not impacted by either the stalling conditions nor the thermal stress. Photosynthetic rates remained stable at about 4 μL.O_2_.mgC^−1^.h^−1^ with no significant changes between all conditions (Fig. 5a, Supplementary Material Table 2) and the maximal photosystem II quantum yield (F_V_/F_M_) average was 0.58 ± 0.04, a nearly optimal value for phytoplanktonic cells (Fig. 5b, Supplementary Material Table 3). No photoprotective processes of energy dissipation were observed (Supplementary material Table S3). At CT-T2 and during the whole heat-stress experiment, the number of observed symbionts per microscopic field decreased, as suggested by the symbiont density counts. Along the experiment, we also observed that the variability of the F_V_/F_M_ increased both within colonies and between colonies (Fig. 5b, Supplementary Material Figure 4).

**Fig. 5:**
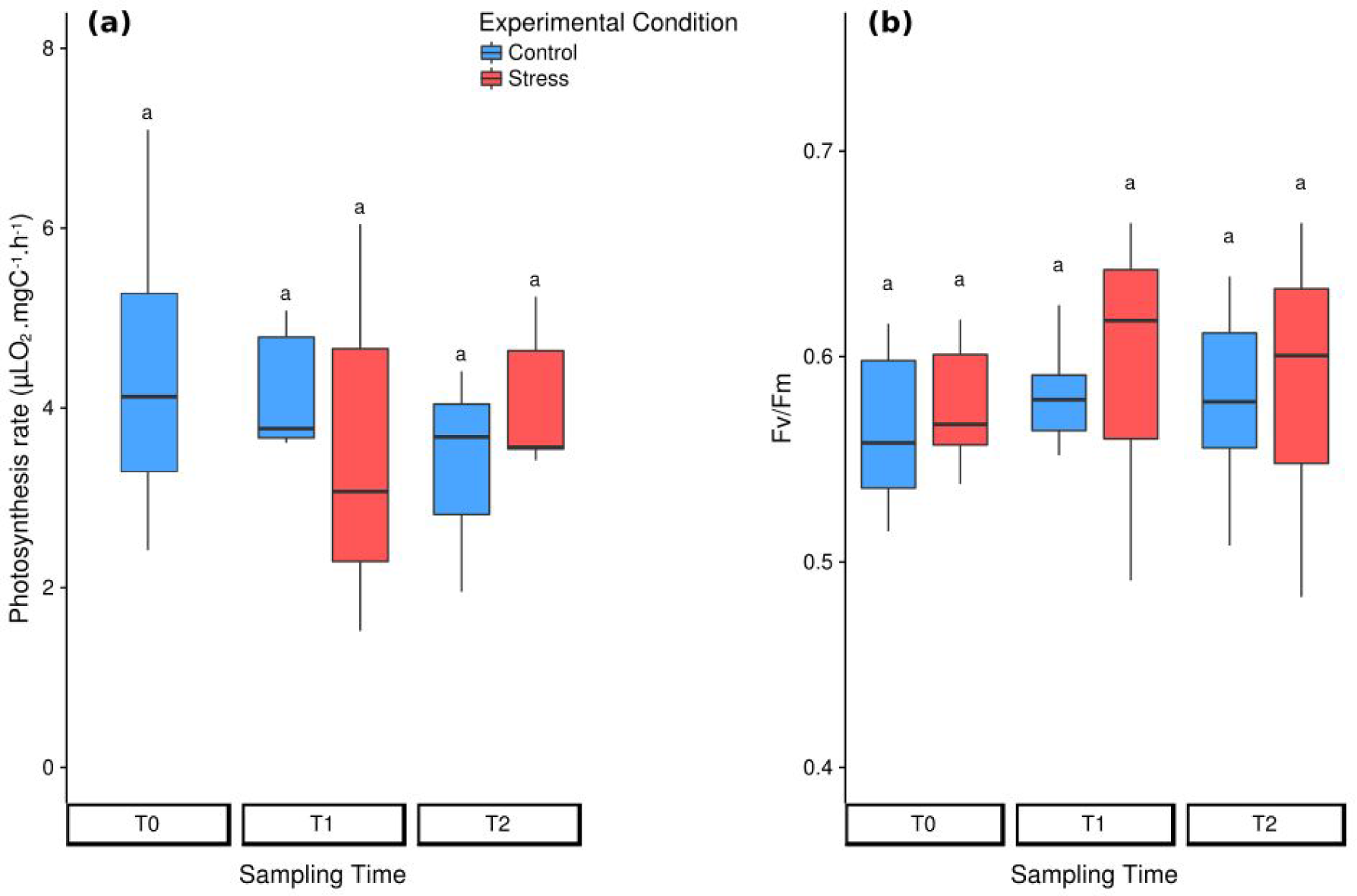
Photosynthetic rates of the symbionts in control (in blue) and heat-stressed (in red) colonies of *Collozoum pelagicum.* **(a)** Box-plots of photosynthesis rates measured by oxygen consumption, when normalized by the symbiont surface of the colony. **(b)** Box-plots of PAM-microscopy measures. Results from 2 way-ANOVA testing Sampling Time, Treatment and Interaction effects are represented by Tukey post-hoc test associated letters for interaction effect.

### Symbionts morphological changes

TEM micrographs were used to monitor ultrastructural changes of the symbiotic *B. nutricula* cells. In control colonies, vacuolar structures containing filamentous particles that could be degradation residues of organelles by virus-like particles (VLPs) were observed in cells at T1 (Fig. 6a, Supplementary Material Figure 5). At T2, these filamentous particles were still present and apoptotic bodies containing membrane-packaged cellular debris appeared as multiple electron-dense bodies within the symbiotic cells (Fig. 6b) but nuclei as well as chloroplast ultrastructure remained preserved (Supplementary Material Figure 6).

**Fig. 6:**
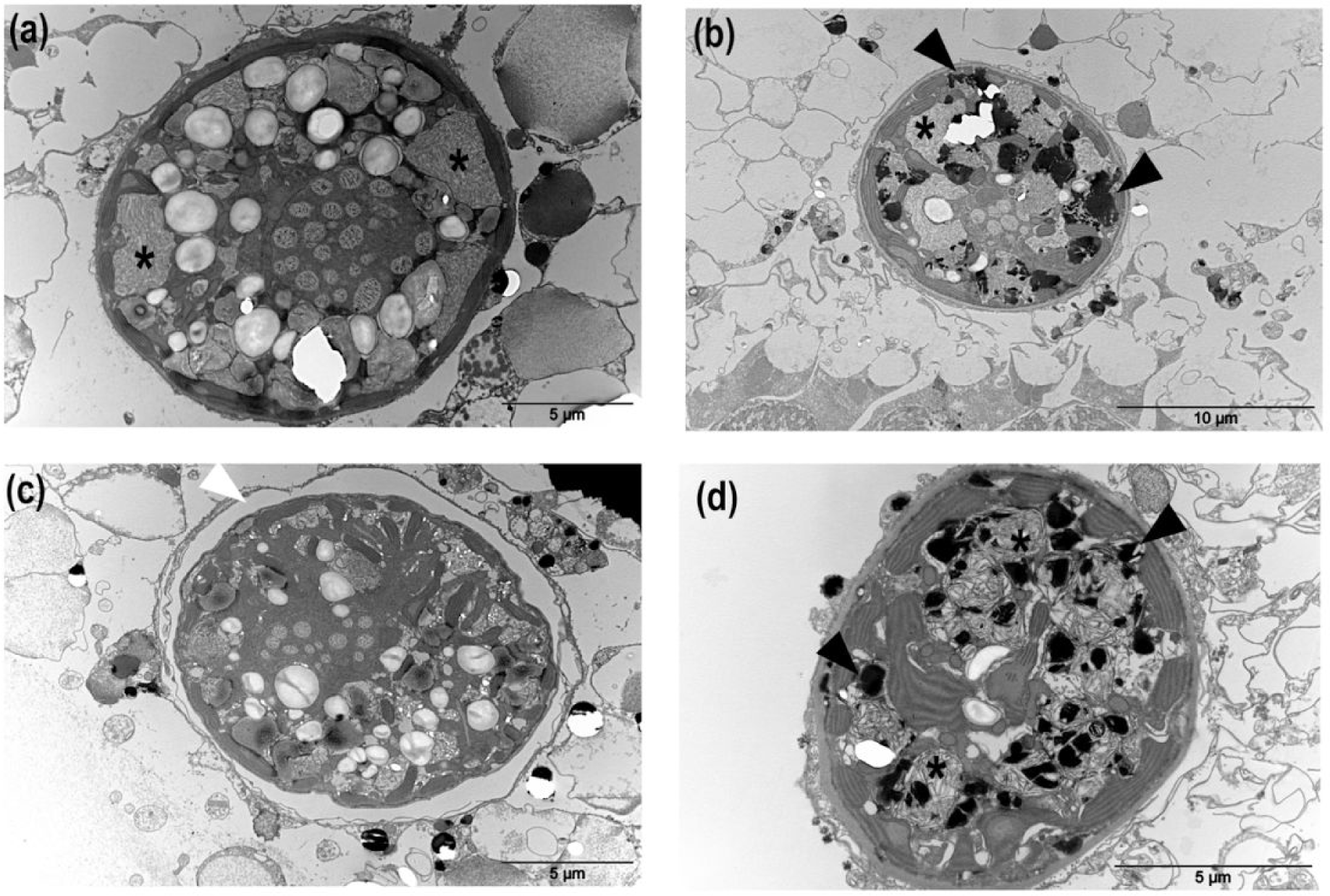
Morphological structures corresponding to the main cellular responses induced in *in hospite Brandtodinium* cells in control and heat-stressed colonies of *Collozoum pelagicum.* Transmission electron microscopy of *Collozoum pelagicum* cross section in control T1 **(a)**, control T2 **(b)** at 21°C, and thermal stress T1 at 25°C **(c)** and thermal stress T2 at 28°C **(d)**. Asterisks: virus-like particles, black triangles: apoptotic bodies, white triangles: vacuolar matrix.

In thermal-stressed colonies, filamentous particles were also observed but to a lesser extent. Instead, an important vacuolization around the symbionts, thylakoid disorganization, nucleus dissolution and destabilized organelles structures were observed at 25°C (Fig. 6c), suggesting autophagic processes. At 28°C, the *Brandtodinium* cells from heat-stressed colonies showed marked necrosis patterns with numerous apoptotic bodies and their cellular content appeared completely shrinked, with abundant electron-dense bodies (Fig. 6d). Inside decaying symbiotic cells, chloroplasts were the only healthy organelles, apparently, consistent with the maintenance of the photosynthetic capacities observed in the physiological study (Fig. 6; Supplementary Material Figure 6).

## Discussion

Bleaching is commonly described in the phylum Cnidaria as a loss of colour originating in the exclusion of the symbiotic *Symbiodinium* dinoflagellates from the animal, and/or the degradation of the photosynthetic pigments within the symbiont chloroplasts (Douglas, 2003). In our experiment, we observed a heat-induced decrease of dinoflagellate density in Collodaria, a planktonic group in symbiosis with the dinoflagellate genus, *Brandtodinium.* It shows that bleaching processes are not be restricted to benthic fauna in symbiosis with *Symbiodinium* dinoflagellates, but likely constitute a common stress response in marine organisms harboring photosymbionts. Similarly to what has been described in cnidarians (Weis, 2008), the decrease of symbiont density in *C. pelagicum* may come from *B. nutricula* digestion or expulsion. It has been shown that, under standard conditions, cnidarians continuously expulse degraded forms of *Symbiodinium* to maintain a healthy population in symbiosis, whereas in stress conditions healthy algal cells are also released in the environment (Fujise *et al.*, 2014). In our study, the number of symbionts located at the vicinity of the central capsules decreased, but symbiont density in the gelatinous matrix is constant. Assuming that expulsion of symbionts requires to go through the gelatinous matrix, an increased density could have been expected . Unfortunately, we were not able to collect *B. nutricula* cells from the seawater surrounding colonies to check their physiological status during the experiment. Still, all symbionts observed inside the hosts by electron microscopy showed signs of PCD-like mechanisms such as autophagy or apoptosis, suggesting cell decay. Apoptosis is triggered by a series of controlled cellular processes involving caspases which cleave key proteins, cytoskeletal elements and cell adhesion molecules (Dunn et al., 2007). In the autophagic pathway, the target is enveloped in a membrane structure before fusing with lysosome, which will engaged digestion by hydrolytic enzymes (Cuervo, 2004). Both apoptosis and autophagy have been observed during cnidarian bleaching and may occur together (Dunn et al., 2007) or alternatively, especially in response to hyperthermal stress (Richier et al., 2006; Dani et al., 2016). Such PCD-like mechanisms have also been described in phytoplankton (Bidle, 2015). In addition, another process related to symbiotic dinoflagellate autophagy has been reported in cnidarians and referred to as “symbiophagy”, which results in the digestion of the resident dinoflagellates (Downs et al, 2009). TEM observations in *C. pelagicum* suggested that a similar diversity of PCD-like mechanisms would be involved in planktonic symbiosis breakdown.

In our study, respiration rate increased between 21 and 25°C. As respiration rates were not measured on the same specimen over the course of the experiment, and only at two different temperatures, this experiment only estimate the theoretical Q10 for Collodarian. The Q10 value of 2,66 for Collodarian falls into the Q10 range of 2 to 3 estimated for zooplankton after compilation of different studies (Hernández-León and Ikeda, 2005; Ikeda 2014). Respiration rate increase as a function of temperature has already been reported in a large panel of zooplankton group such as Rhizaria, Copepods, Tintinnids (Lombard et al., 2005; Vidal, 1980; Verity, 1985) and could be due to several processes. The global rates of all chemical reactions increase with temperature (Arrhenius 1889) and, until enzymatic structure is altered, most biological enzymatic processes require oxygen, so the oxygen demand increases with temperature. Also, higher temperatures likely enhance the growth of prokaryotes associated to the host and in turn contribute to a not negligible part of the holobiont respiration (del Giorgio et al., 1997). Finally, it has also been shown that digestion increases the respiration rate of organisms (Conover, 1978).

Virus-like particles could also be involved in symbiont degradation as we observed features similar to the viruses described as spreading throughout *Symbiodinium* cells (Weynberg et al. 2017). Several cases of nucleocytoplasmic large DNA and single stranded RNA viruses infecting *Symbiodinium* have been reported (Correa et al., 2016; Weynberg et al., 2017). Such lysis of symbiotic dinoflagellates by viruses supports the microbial bleaching hypothesis, which suggests that bleaching could be initiated by the microbial community shift induced by heat-stress (Thurber et al., 2017). Viral infections in marine algae have been shown to promote ROS activity, triggering subsequent caspase activity and further programmed cell death in host cells (Bidle & Vardi, 2011; Bidle 2015). Recently, Weynberg et al. (2017) proposed that the cnidarian endosymbiont *Symbiodinium* harbors a lysogenic virus within its genome that would induce a lytic infection cycle under stressed conditions. These authors observed that if nuclei structures were rapidly degraded by virus infection, other organelles like chloroplasts and mitochondria remained almost intact until infectious particles filled the entire cell structure. In our experiment, we also showed that chloroplasts remained morphologically intact even though other *B. nutricula* organites were severely damaged. This observation is consistent with the relatively constant and high photosynthetic efficiency measured throughout the experiment. To our knowledge, only one study has described large icosahedral virus-like particles associated with vacuolar structures in phaeodarians, a taxonomic group close to radiolarians, within the Rhizaria lineage (Gowing, 1993). This study suggests that VLPs could have been acquired by the host through feeding on sinking particles, while in our case, it seems that the viral attack was restricted to the symbiont.

Based on our observations (e.g. respiration virtually stopped, morphological structure altered), a temperature of 28°C *(i.e.* T2) likely exceeded the *C. pelagicum* colonies tolerance threshold. The pronounced thermal stress we applied in our experiment aimed at observing and characterizing a marked response of the symbiotic system, likely preventing the potentially existing acclimation period required for the host plastic response. The free-living stage of the microalgal symbionts *B. nutricula* tolerate temperatures up to 29°C in culture experiments, displaying relatively high growth rates and photosynthetic efficiency (Supplementary Text 1 & Supplementary Material Figure 7). *Symbiodinium* physiology was also more affected by heat stress when in symbiosis with corals than when maintained free-living in culture (Buxton et al., 2009). Photosymbiosis induces critical symbiont morphological and physiological changes (i.e. loss of motility) likely leading to a greater sensitivity when host protection is altered and prevent buffering external stressors.

## Conclusions

In this study, we show that collodarians are likely sensitive to heat-induced bleaching. We unveil several features of the processes involved, such as host cells degradation, partial digestion and potential viral infection of the symbionts, while the chloroplast is clearly the ultimate structure to remain active. These results raise questions such as regarding underlying triggering and cellular processes involved in symbiont loss in *C. pelagicum* or preservation mechanisms of chloroplasts during symbiont degradation. Further analysis using complementary approaches like transcriptomic or metabolomic would certainly be helpful to decipher more precisely the holobiont response to elevated temperatures.

## Supplementary data

Text S1-S2: Provided separately as .pdf

Figure S1-S7: Provided separately as .pdf

Table S1-S3: Provided separately as .xlsx

## Acknowledgements

The authors greatly acknowledge the Centre Commun de Microscopie Appliquée (Université Côte d’Azur), especially Sophie Pagnotta who performed TEM image acquisitions. The authors acknowledge the members of Villefranche-sur-mer oceanological observatory for their help during the experimental work, especially Simon Ramondenc, Guillaume de Liege, David Luquet, Sophie Marro and Régis Lasbleiz.

## Funding

This work benefited from the support of the projects IMPEKAB ANR-15-CE02-0011 and inSIDE ANR-12-JSV7-0009-01 of the French National Research Agency (ANR). This research was supported by the research infrastructure EMBRC-Fr (www.embrc-france.fr).

## Author Contribution

EV, MMS, CB, EB, TL, FN performed the experiment. EV performed the PAM measures, image analysis and the statistical treatments and wrote the manuscript. VD contributed to TEM analysis and manuscript writing. MMS performed the oxygen measurements. CB performed the sequence analysis. CB, FL, CS, CS and FN commented and contributed to the final version of the manuscript. FN planned and designed the research.

